# Tracking cytosine depletion in SARS-CoV-2

**DOI:** 10.1101/2020.10.26.354787

**Authors:** Ruibang Luo, Yat-Sing Wong, Tak-Wah Lam

## Abstract

**Motivation:** Danchin et al. have pointed out that cytosine drives the evolution of SARS-CoV-2. A depletion of cytosine might lead to the attenuation of SARS-CoV-2.

**Results:** We built a website to track the composition change of mono-, di-, and tri-nucleotide of SARS-CoV-2 over time. The website downloads new strains available from GISAID and updates its results daily. Our analysis suggests that the composition of cytosine in coronaviruses is related to their reported mortality. Using 137,315 SARS-CoV-2 strains collected in ten months, we observed cytosine depletion at a rate of about one cytosine loss per month from the whole genome.

**Availability:** The website is available at http://www.bio8.cs.hku.hk/sarscov2/.

**Supplementary information:** Supplementary data are available at *Bioinformatics* online.

## 1 Introduction

SARS-CoV-2 is a positive-strand RNA virus. The intracellular multiplication of positive-strand RNA viruses requires complex interactions with the host cell. The coronavirus genome mimics the structure of a cellular mRNA. It is rapidly translated into an RNA-dependent replicase and proteins that allow the virus to hijack specific host functions used during viral translation, RNA replication, and viral envelope construction. The virus uses a critical set of metabolic pathways to access the pool of ribonucleotide triphosphates needed for the transcription of many copies of the replicated RNA strand. This makes the viral sequence highly sensitive to changes in the nucleotide pool, the composition of which might be reflected in the virus evolution as it mutates. Although coronaviruses have evolved a specific function meant to overcome some of this limitation via a proofreading step in the replicase function, yet mutations remain unavoidable and viruses, which generate a vast number of particles within a single patient, will evolve quasi-species that will progressively integrate the various types of selection pressure that the virus faces.

Among the pressures, the availability of CTP (Cytidine triphosphate, and hence of cytosine-based precursors) might be an important driving force in the evolution of the virus. The CTP supply in a human is lower than the three other essential triphosphates ATP, GTP, and TTP for RNA synthesis, and the scarceness of CTP in the host will, in turn, lowers the composition of cytosine in a virus genome along with its evolution (Armengaud, et al., 2020; Danchin and Marliere, 2020). This phenomenon is called depletion, and it co-occurs with the decreased virulence of an RNA virus commonly observed in an outbreak. If under constant selection pressures, the speed of depletion tends to stabilize, but if extraneous pressures such as massive vaccination or effective medication, the depletion may slow down or even reverse (Danchin and Timmis, 2020). By evaluating the speed of cytosine depletion in the new SARS-CoV-2 strains, it might 1) be used as a new metric at the molecular level for gauging the speed of the spread of SARS-CoV-2, and 2) predict the resistance of SARS-CoV-2 to new vaccines and drugs.

## 2 Methods

We built an interactive website at http://www.bio8.cs.hku.hk/sarscov2 to show the mono-, di-, and tri-nucleotide composition trends of the whole genome and single genes. The SARS-CoV-2 genomes used in this website are from GISAID (https://www.gisaid.org/). The acknowledge table of the strands used in Table 1 is at http://www.bio8.cs.hku.hk/sarscov2/gisaid_cov2020_76.xls.

**Table 1:**
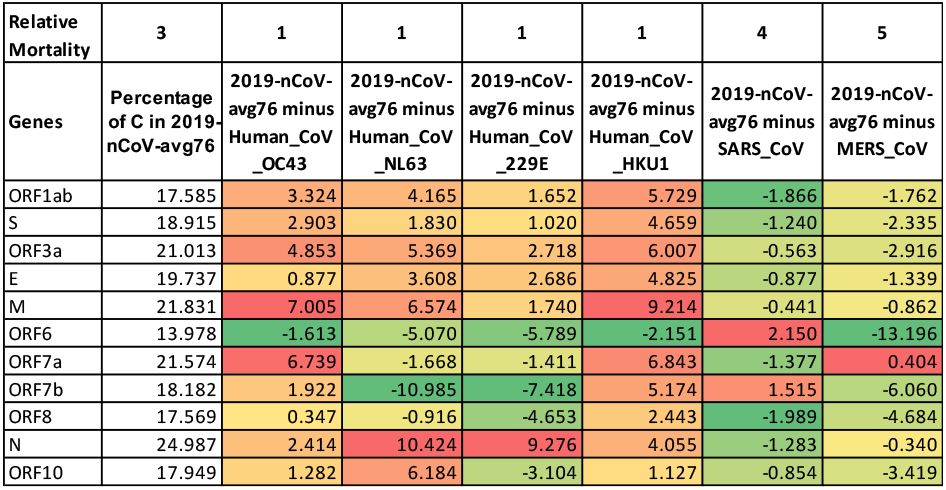
The relative composition of cytosine in the eleven genes (using the annotation of Wuhan-Hu-1) of four common cold coronaviruses, SARS and MERS comparing to the average of 76 SARS-CoV-2 strains (2019-nCoV-avg76, collected from Dec 20, 2019 to Feb 15, 2020). The mortality levels are set according to their reported mortality.

New SARS-CoV-2 strains are retrieved from the GISAID platform on a daily basis. We used only full genomes and removed those sequences short than 29kbp after removing gaps (runs of ‘N’s). IUPAC bases are substituted with C or the alphabetically smaller base. We used “BetaCoV/Wuhan/WIV04/2019” (https://www.ncbi.nlm.nih.gov/nuccore/MN996528) and its annotations as our reference. For every full genome, use aligned it against the reference using MAFFT (Katoh and Standley, 2013). The completeness at both ends of the full genomes varied. Thus, we removed the bases aligned with the head 130bp and tail 15bp of the reference.

## 3 Results

In **Table 1**, we first compared the composition of nucleotides in multiple coronaviruses, including SARS-CoV-2 (an average of 76 strands collected from Dec 20, 2019 to Feb 15, 2020), SARS, MERS, and four that causes a common cold. Compared to SARS-CoV-2, which we assumed a relative mortality level of 3, the percentage of cytosine is lower in almost all of the eleven genes in the four common cold coronaviruses with a lower mortality level 1. However, the percentage of cytosine is higher in SARS (mortality level 4) and MERS (mortality level 5). The result suggests that cytosine depletion is associated with reduced morbidity of coronaviruses.

Next, we summarized the trend of different nucleotides in SARS-CoV-2 strains collected from Dec 20, 2019 to Oct 2, 2020. The results are available on an interactive website at http://www.bio8.cs.hku.hk/sarscov2. 137,315 strains from GISAID (Elbe and Buckland-Merrett, 2017) were used (see Methods). The trend of mono-nucleotides A, C, G, and T of the whole genome and ten genes are shown in **Figure 1** (whole genome, orf1ab, S, E, M, and N), and **Supplementary Figure 1** (whole genome, NS3, NS6, NS7a, NS7b, and NS8). The whole genome shows a sign of cytosine depletion, at a rate of about one cytosine loss per month. Except for gene E, N, NS7a, and NS7b, all other genes show a sign of cytosine depletion. The trend of di-nucleotides and tri-nucleotides is available on the website. Looking into the trend of di-nucleotides in the whole genome, all di-nucleotides containing a cytosine are having a sign of depletion except for “GC”.

**Figure 1:**
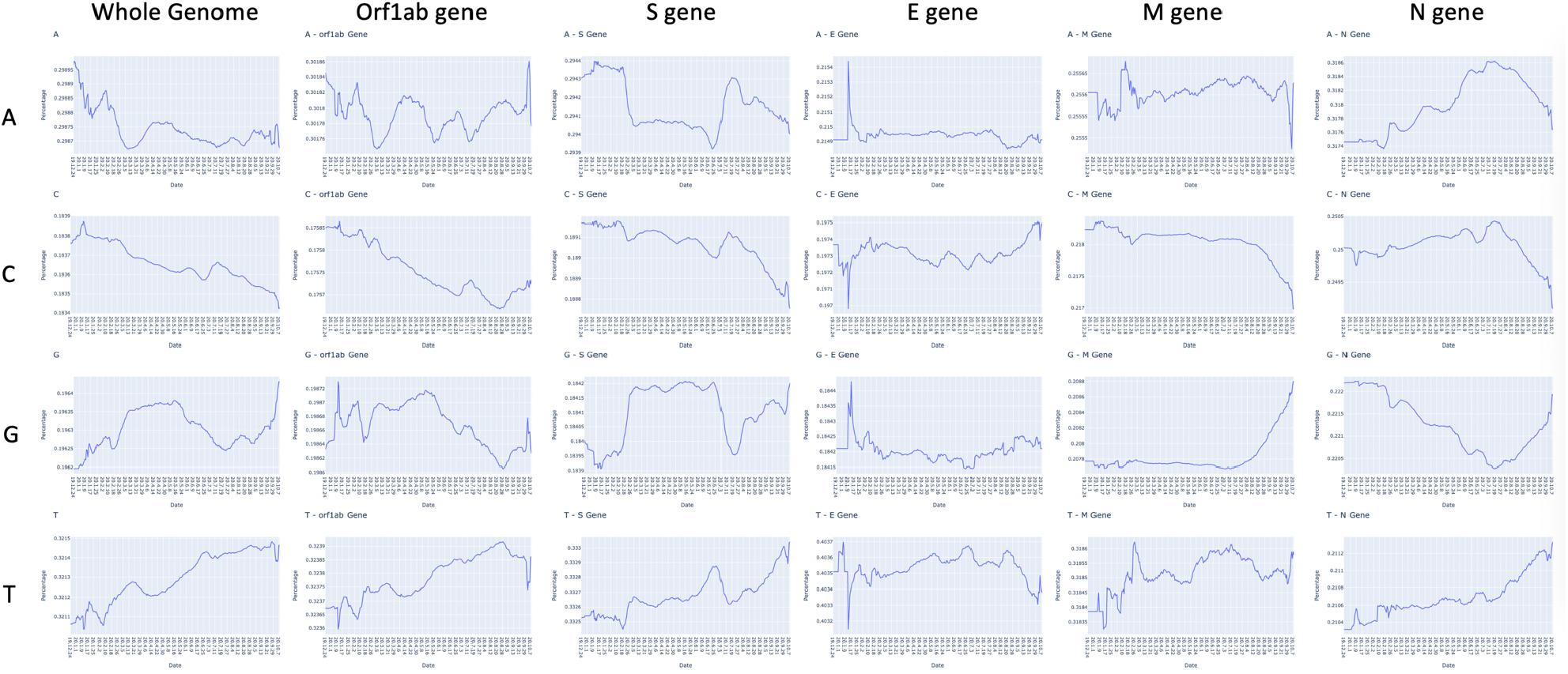
The trend of mononucleotide A, C, G and T in 137,315 SARS-CoV-2 strains collected from Dec 20, 2019 to Oct 2, 2020. The data points were smoothed by a 15-day window.

## References

Armengaud, J., et al. The importance of naturally attenuated SARS-CoV-2 in the fight against COVID-19. Environ Microbiol 2020;22(6):1997–2000.

Danchin, A. and Marliere, P. Cytosine drives evolution of SARS-CoV-2. Environ Microbiol 2020;22(6):1977–1985.

Danchin, A. and Timmis, K. SARS-CoV-2 variants: Relevance for symptom granularity, epidemiology, immunity (herd, vaccines), virus origin and containment? Environ Microbiol 2020;22(6):2001–2006.

Elbe, S. and Buckland-Merrett, G. Data, disease and diplomacy: GISAID’s innovative contribution to global health. Glob Chall 2017;1(1):33–46.

Katoh, K. and Standley, D.M. MAFFT multiple sequence alignment software version 7: improvements in performance and usability. Mol Biol Evol 2013;30(4):772–780.

