## Supplementary Figure 1 for "Tracking cytosine depletion in SARS-CoV-2"

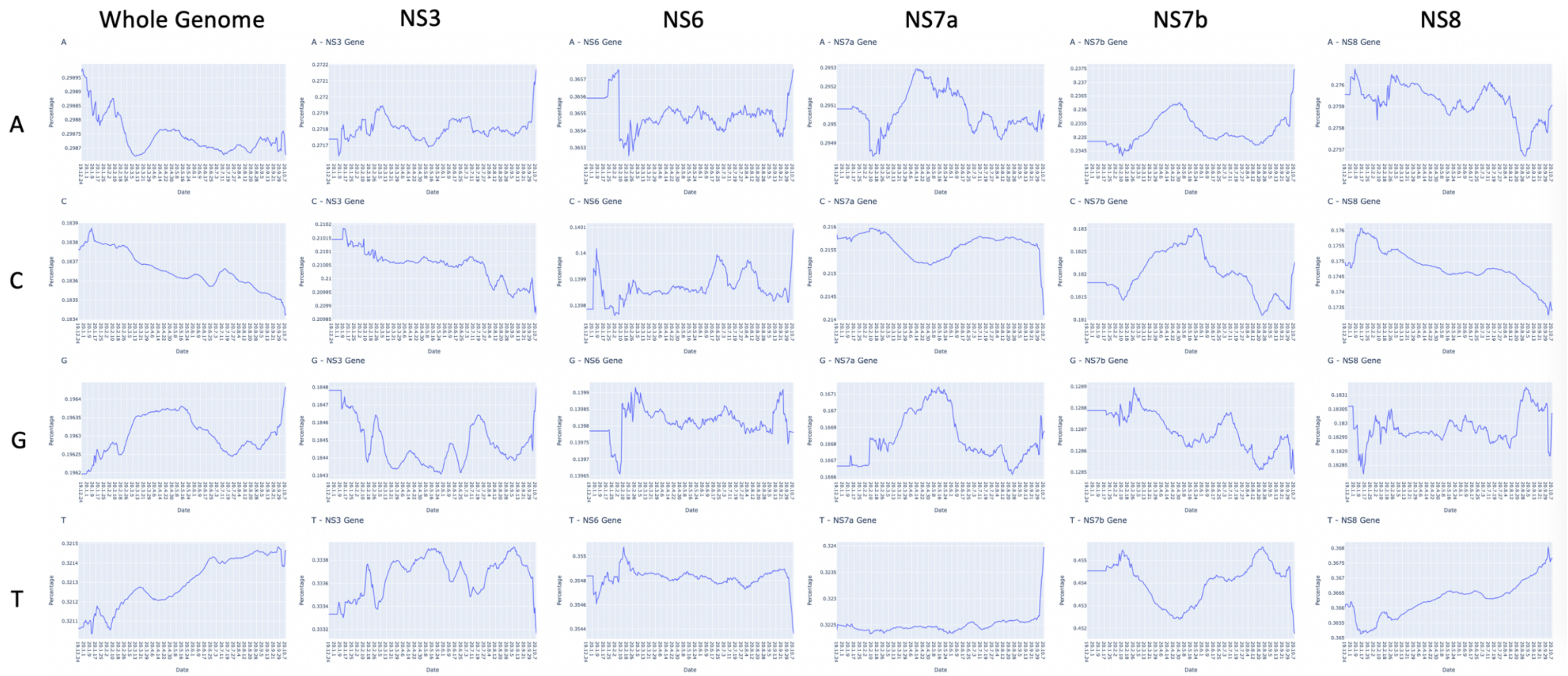


**Supplementary Figure 1:** The trend of mononucleotide A, C, G and T in 137,315 SARS-CoV-2 strains collected from Dec 20, 2019 to Oct 2, 2020. The data points were smoothed by a 15-day window.
